# Belief loads of assumptions impact brain networks underlying logical reasoning: A machine learning approach

**DOI:** 10.1101/2020.05.16.092304

**Authors:** Maryam Ziaei, Mohammad Reza Bonyadi, David Reutens

## Abstract

Prior knowledge and beliefs influence our reasoning in daily life and may lead us to draw unwarranted conclusions with undesirable consequences. The underlying neural correlates of the interaction between belief and logic, prior to making logical decisions, are largely unknown. In this study, we aimed to identify brain regions important in distinguishing belief load of assumptions in logical decision making. Thirty-one healthy volunteers (18-29 years old) participated in an fMRI study and were asked to respond to a series of syllogistic arguments in which assumptions were either congruent (believable) or incongruent (unbelievable) with common knowledge. An interpretable machine learning algorithm, an L1 regularized Support Vector Machine, was used to explain the discriminatory pattern of conditions given the brain activation patterns. Behavioral results confirmed that believable premises were incorrectly endorsed more than unbelievable ones. Imaging results revealed that several connectivity patterns anchored around the insula, amygdala, and IFG were important in distinguishing believable from unbelievable assumptions at different time points preceding logical decisions. Our convergent behavioral and imaging results underscore the importance of the belief loads of our *assumptions* for a logically sound decision. Our results provide new insights into neural and potential cognitive mechanisms underlying the interaction between belief and logic systems, with important practical implications for social, complex decisions.

**Highlights:**

- Belief load of premises impact logical decisions
- Regional importance in distinguishing belief load changes in different TRs
- Belief load of assumptions elicits emotional and salience responses
- Regional connectivity changes as the reasoning process evolves at different TRs
- Insula, caudate, amygdala, and IFG were among highly connected hubs during the task

**Significance:** It has been experimentally shown that decision-makers often ignore given assumptions in favor of their own beliefs, potentially leading them towards a subjective rather than a logical decision. Consider “Carbon emission tax” given the assumption of “global warming”. If a decision-maker does not believe in “global warming”, the final decision on “Carbon emission tax” is not driven by factual premises but by the personal belief of the decision-maker. Understanding neural mechanisms underlying the interaction between the decision-makers’ beliefs and factual premises sheds light on factors driving belief bias and potential interventions to circumvent it. The main contribution of this study is to investigate neural mechanisms in a logical reasoning task in which the belief load of the assumptions was manipulated.

## Introduction

Logical reasoning requires the recruitment of related facts, assumptions, and prior knowledge to come to reasonable conclusions. Beliefs and prior knowledge, however, may overshadow logic and lead to unwarranted conclusions, a phenomenon known as the belief bias (De Neys, 2012; 2013). Consider, for example, “carbon emission tax” given the assumption of “global warming”; if a decision-maker does not believe in “global warming”, even though supported by scientific evidence, the final decision on “carbon emission tax” might not be driven by the factual premises but by the personal belief of the decision-maker. Syllogistic reasoning, drawing a conclusion from given assumptions, with manipulated belief load is frequently used to study the interaction between belief and logic. To investigate belief bias using syllogistic reasoning, most previous neuroimaging studies have focused on the influence of belief load during the conclusion (decision making) stage (Goel et al., 2004a; Knauff et al., 2003; Monti et al., 2007; Reverberi et al., 2012a). These studies argue that the believability of the conclusion influences reasoning by motivating people to reason more thoroughly when the statements are unbelievable (Evans et al., 1983; Johnson-Laird, 2001).

A few studies, however, have investigated how the belief load of assumptions may impact the performance of decision-makers (Halford et al., 2015; Maybery et al., 1986; Thompson, 1996). These studies have shown that conclusions supported by believable premises are more likely to be accepted, emphasizing the importance of the belief load of the assumptions in drawing correct conclusions. To our knowledge, only one neuroimaging study has measured neural responses in a logical reasoning task when the content of the assumptions or conclusion was not congruent with prior beliefs, known as belief-content conflict (Stollstorff et al., 2012). It was found that the activity in the right lateral prefrontal cortex (PFC) varies with the degree of belief-content conflict, i.e., the number of unbelievable statements in the syllogistic task. The role of other brain areas and their connections underlying the interaction between the decision-makers’ beliefs and factual premises, however, was not thoroughly investigated. Such investigation of neural mechanisms would shed light on factors driving belief bias and potential interventions to circumvent it.

The primary aim of this study was thus to investigate how the neural mechanisms involved in decision making differ when the belief load of given assumptions in a reasoning task is manipulated. Healthy younger adults completed a syllogistic reasoning task in which they were asked to either endorse or reject a conclusion statement, and the belief load of assumptions was either congruent (all parrots are birds) or incongruent (all parrots are lizards) with common knowledge. Based on previous behavioral studies, we expected that participants to be “deceived” by the belief load of the premise, hence, incorrectly endorsing statements with believable premises more often than statements with unbelievable premises. This would consequently lead to a higher false-negative rate and lower false-positive rate in statements with unbelievable premises comparing to believable premises. For the imaging data, we used an interpretable classification method to identify brain regions, characterized by the mean brain activation signals from anatomically defined regions of interest, important in differentiating experimental conditions, defined according to whether participants were processing believable or unbelievable assumptions. This approach reduces the number of features fed into the classifier. Hence, the method does not suffer from the ‘curse of dimensionality’, a common issue in Multivoxel Pattern Analysis (MVPA) algorithms. Based on the probabilistic association between regions, the method was extended to establish a graphical model of the regions that are most likely to work together to explain the experimental condition. Given the complexity of the interactions between belief and logic systems, we anticipated multiple cognitive processes to be involved in differentiating believable from unbelievable assumptions. For instance, we expected that assumptions would involve the retrieval of currently-held beliefs from memory, engaging the hippocampus (Goel et al., 2004b; Ziaei et al., 2020a), and that conflict between assumptions and an individual’s belief is likely to elicit an emotional response, involving the amygdala (Eimontaite et al., 2019; Stollstorff et al., 2012). Inhibition of currently-held beliefs to achieve a logical decision is expected to involve the insula, a critical node of the salience network (Menon, 2015), and the inferior frontal gyrus, a primary node in inhibitory control (Aron et al., 2014; Ziaei et al., 2020b).

## Materials and Method

### Participants

Thirty-one healthy young adults participated in this study. Two participants’ imaging data were excluded due to technical issues such as failure of the image reconstruction stage, extensive head movement during scanning, and corruption of data. Therefore, only twentyeight participants were included in the final analyses (age 18-26 years, *M* = 21.13, *SD* = 2.72; 15 females). All participants were students recruited from the University of Queensland in exchange for course credits or a payment of $20 per hour. Screening for claustrophobia, neurological and psychiatric disorders, and magnetic resonance imaging (MRI) incompatibility was conducted prior to the experiment. All participants were right-handed English speakers, had normal or corrected-to-normal vision, and had no history of neurological impairment or psychiatric illnesses. They first undertook the fMRI scanning task, followed by a break, and then completed the neuropsychological assessments. Participants provided written consent and were debriefed upon completion of the experiment. The experiment was approved by the Bellberry Human Research Ethics Committee.

### Materials

In brief, logical arguments used in this study included three statements, two premises and one conclusion, in the form of a standard syllogism. In a premise, the subject and the predicate are arbitrary sets (e.g., dogs, mammals, furniture). Exactly one set is *shared* between the two premises that may appear in either the subject or the predicate in either of the premises. Hence, an argument with two premises involves exactly three sets (*Set*_1_, *Set*_2_, and *Set*_3_ in the example), two uniquely used in each premise and one used in both. The conclusion of a syllogism is a statement about the sets that appear uniquely in premises (*Set*_1_ and *Set*_3_ in the example). A conclusion “follows” from the premises if the premises provide conclusive evidence to support it. Otherwise, the conclusion “does not follow” from the premises. This includes a conclusion that is wrong, given the premises, or is not completely supported by the premises.

The conclusion and the premises can be believable or unbelievable statements. For example, “all birds are mammals” is an unbelievable statement while “all parrots are birds” is a believable statement. In addition, premises were used to provide a neutral context as a control, for example, “all dogs are sothods”, where “sothods” is a pseudo-word. This type of statement was only used in the premises in a way that the pseudo-word was always the shared set between the premises. Hence, the premises were either believable, unbelievable, or neutral while the conclusion was either believable or unbelievable. The phrases “believable premises”, “neutral premises”, and “unbelievable premises” refer to the believability load of the second assumption. The following are two examples of syllogisms with a believable premise / unbelievable conclusion and an unbelievable premise / believable conclusion respectively:

All sparrows are birds; all birds are animals; therefore, no animals are sparrows (believable premise/unbelievable conclusion)

All oranges are citrus; no citrus are fruits; therefore, some fruits are not oranges (unbelievable premise/believable conclusion)

In a syllogism, in addition to a subject, a predicate and a proposition, each statement includes a *quantifier (All, No, and Some*) and a *copula* (*Is or Is not*). Two types of *proposition* were used, defined by the following combinations of <quantifier, copula>:

1. Proposition type *A*: <All, is>,
2. Proposition type *E:* <No, is>

From here on, we denote syllogisms as <premise 1; premise 2; conclusion> for simplicity. In all of our premises, we used proposition types *A* and *E* (counterbalanced across all runs). We avoided the use of proposition *E* in both premises as it simplifies the reasoning task. For the conclusions, however, we used the propositions required to ensure that conclusions that followed the premises and those that did not follow the premises were balanced. We controlled for the difficulty and negation in all syllogisms.

A total of 96 syllogisms was generated, comprising a total of six conditions: 1. believable premise / believable conclusion, 2. believable premise / unbelievable conclusion, 3. unbelievable premise / believable conclusion, 4. unbelievable premise / unbelievable conclusion, 5. neutral premise / believable conclusion, 6. neutral premise / unbelievable conclusion. An algorithm that creates all the syllogisms based on criteria specified above was developed (see Supplementary Material for access to all the syllogisms used in the study).

### Task design

The scanning session lasted 45 minutes and included two components: structural MRI (sMRI) and the logical reasoning task with fMRI. Prior to the scan, participants were verbally instructed about the task and a practice run was continued until they were familiar with the timing and instruction of the task. During the logical reasoning task, participants were instructed to ascertain if the conclusion logically followed from the premises. They responded with two keys on an MRI-compatible response box. The first premise was presented for 2 seconds followed by a second premise for 4 seconds. After the second premise, the conclusion statement was presented for 12 seconds. To minimize the working memory load, all statements remained on the screen until the end of the presentation of the conclusion. A fixation cross was presented after the conclusion which was randomly jittered using four intervals: 0.5 seconds (24 trials), 1 second (24 trials), 1.5 seconds (24 trials), and 2 seconds (24 trails) across all runs. The task consisted of 6 runs, each run lasting for 5.16 minutes. Three runs of the task were presented before the sMRI and three after the sMRI.

### Background measures

After a break following completion of the imaging session, all participants were asked to complete a range of background measures assessing executive control such as a Stroop task (Jensen et al., 1966) and Trail Making Test (Reitan et al., 1986) and intelligence measured by National Adult Reading Test (Nelson, 1982); emotional well-being measured by Depression, Anxiety, Stress Scale (DASS-21, Lovibond & Lovibond, 1995^25^). Descriptors of background measures are reported in Table 1.

**Table 1.** Descriptive statistics of logical reasoning task performance and background measures

| *Measure* | *Conditions* | *Mean (Standard Deviation)* |
| --- | --- | --- |
| **Age** |  | 20.65 (2.66) |
| **Gender** |  | 15 females, 14 males |
| **Education (years)** |  | 15.28 (1.97) |
| **DASS – 21** |  |  |
| *Stress* |  | 6.06 (5.19) |
| *Anxiety* |  | 2.68 (3.55) |
| *Depression* |  | 3.31 (4.60) |
| **NART FSIQ** |  | 112.02 (4.77) |
| **Stroop Test in second** |  |  |
| *Congruent* |  | 0.72 (0.12) |
| *Incongruent* |  | 0.82 (0.16) |
| *Neutral* |  | 0.70 (0.10) |
| *Stroop effect* |  | 0.16 (0.12) |
| **Premise condition** |  |  |
| *RT (sec)* | *Unbelievable* | 4.81 (1.37) |
|  | *Believable* | 4.70 (1.22) |
|  | *Neutral* | 4.98 (1.20) |
| *Rejection rate* | *Unbelievable* | 0.98 (0.13) |
|  | *Believable* | 0.93 (0.07) |
|  | *Neutral* | 0.97 (0.13) |

*Note.* NART FSIQ = National Adult Reading Test Full-Scale Intelligence Quotient; Stroop effect = (Incongruent – neutral/neutral); DASS-21 = Depression, Anxiety, Stress Scale with 21 items; Rejection rate = number of times each participant correctly rejected a given syllogism divided by the total number of statements that should have been correctly rejected; SD = Standard Deviation; Stroop effect = (Incongruent – neutral/neutral) \*100; RT = reaction time; sec = second; M = mean; SD = standard deviation.

### Image acquisition

Functional images were acquired at the Centre for Advanced Imaging using a 3T Siemens scanner with a 32-channel head coil. The functional images were obtained using a wholehead T2*-weighted multiband sequence (473 interleaved slices, repetition time (TR) = 655ms, echo time (TE) = 30ms, flip angle = 60°, field of view (FOV) = 190mm, multi-band acceleration factor = 4, voxel size = 2.5mm^3^). High-resolution T1-weighted images were acquired with an MP2RAGE sequence (176 slices with 1mm thickness, TR = 4000ms, TE = 2.89ms, TI = 700ms, FOV = 256ms, voxel size = 1mm^3^). Participants observed the tasks on a computer screen through a mirror mounted on top of the head coil. To reduce the noise and minimize head movement, participants were provided with earplugs and cushions inside the head coil.

### Preprocessing

For functional analysis, T2*-weighted images were pre-processed with Statistical Parametric Mapping Software (SPM12; http://www.fil.ion.ucl.ac.uk/spm) implemented in MATLAB 2015b (Mathworks Inc., MA). Following the realignment to a mean image for head-motion correction, images were segmented into gray and white matter. Then, images were spatially normalized into a standard stereotaxic space with a voxel size of 2 mm^3^, using the Montreal Neurological Institute (MNI) template, and then spatially smoothed with a 6 mm Gaussian Kernel.

### Signal extraction (MarsBar settings)

For each condition, participant, and repetition time during the conclusion stage, we extracted the mean BOLD parameter estimate value using the MarsBar toolbox (order 2 autoregressive model) for 50 hand-picked ROIs, taken from the AAL (anatomical automatic labelling) atlas (Tzourio-Mazoyer et al., 2002).

### Data analysis

Our aim was to investigate whether the believability of premises would lead to a significant difference in brain activation during the conclusion stage. For this reason, we only included believable and unbelievable conditions and compared the brain signals for each regions of interest in these two conditions only. We first extracted signals for 50 handpicked *a priori* regions at each TR during the conclusion stage (18 TR) for both believable and unbelievable premise conditions. This process provided 2 × 28 samples (56 samples in total with 2 conditions and 28 individuals) for each TR, with each sample involving a mean value for 50 regions of interests. The selection of regions was based on their involvement in cognitive functions deemed to be important based on previous literature and meta-analysis in logical reasoning (Prado et al., 2011). These regions included areas from prefrontal and parietal lobes, as well as subcortical areas such as the amygdala, hippocampus, caudate and putamen. As the number of picked regions used as features is smaller than the sample size (56), our method does not suffer from the issue of large feature size and small sample size.

#### Separating conditions at each TR

At each TR, the samples formed independent variables for an interpretable classification method, specifically the support vector machine (SVM) with L_1_ regularization (Bi et al., 2003), to separate believable and unbelievable premises, labelled as —1 and 1. Interpretable classifiers such as SVM not only distinguish between classes but also provide insights to the importance of features (regions) involved in the classification (Pereira et al., 2009; Tibshirani, 1996; Weston et al., 2001). Therefore, an interpretable method can be used to rank features according to their importance in distinguishing between experimental conditions.

At each TR, we performed a shuffle test to determine if the accuracy of the classifier in separating the conditions was significantly better than chance (the null hypothesis was that the premise load could be described by the regions’ activity at the chance level, *t-test*, *p*<0.05) (Pereira et al., 2009). Specifically, we ran the classifier 2000 times and, for each run, we selected a subset of participants (90% of participants, standardized to mean equal to zero, standard deviation equal to one) as a training set. We picked the regularization parameter for SVM-L1 using a 5-fold cross validation on the range of [0, 1] with a step size of 0.1 for the training set. We then trained the classifier using this regularization factor on the training set and calculated the accuracy using the remaining participants as the test set (10% of participants, transformed using the standardization factors obtained for the training set). We used the area under the curve (AUC) of the receiver operating characteristic (ROC) curve to measure accuracy. Then, on the same training set, we shuffled the labels randomly, found the best regularization factor, trained the classifier, and calculated its accuracy on the test set using a similar standardization procedure to the non-shuffled case. A *t-test* was used to compare the performance of the classifier trained on the original training set and the shuffled training sets, the null hypothesis being that the classifier performs at chance level (Pereira et al., 2009). To decrease the population biases and impact of outliers on our analyses, we implemented a bootstrap aggregation (bagging) method (Jollans et al., 2019), as described below.

#### Stability of regional contributions to discriminative pattern of conditions at each TR

SVM-L_1_ provides a hyperplane with sparse coefficients (Bi et al., 2003). The sparsity of the coefficients enables their use as a measure of the importance of features (in this case the importance of the regions) in separating the conditions. This direct approach, however, is prone to population effect and outliers (Meinshausen et al., 2010). To address this issue, we used a bootstrap aggregation (bagging) method to find a stable set of regions that contribute to the discriminative pattern (Rondina et al., 2013). We ran the SVM_L1 2000 times on a subset of participants and regions (50% of regions, called *in-bag* regions) selected randomly at each run. The distribution of the coefficient values of each region across runs provides an insight into the importance of that region. The fraction of the runs in which a region was selected for training and had a non-zero coefficient value after training was called the *importance stability factor* (ISL). The ISL indicates the stability of the contribution of the region in separating the classes (the larger the ISL, the greater the stability). Defining *r_i_* the event of a region *i* being important, the ISL of that region provides the probability of that region being important, denoted as *P*(*r_i_*). This procedure reduces the dependence of estimating regional importance on the population (Meinshausen et al., 2010; Rondina et al., 2014; Schrouff et al., 2018).

The absolute value of the coefficients found by SVM-L1 provides an insight into the importance of the region in separating the conditions, for samples standardized in the same manner as for the training set. For coefficients that were flagged as stably important using the above procedure, we calculated the median of the absolute value of the coefficient and its 95% confidence interval using a bootstrap aggregation method (5000 resampling for calculation of the mean). This median, called the *bagged importance level* (BIL) throughout the paper, for each stably important region was used to rank the regions in terms of their importance in separating the conditions.

#### Probability of functional connections between regions to explain the response patterns

Under the *i.i.d* assumption, the non-zero coefficients of SVM-L1 indicate a sparse network of regions functioning together to explain the response pattern. This network, however, may vary for each run (bag) when the bagging approach is used because the instances and features involved at each run may be different from one another. Therefore, we modelled the bagging process by a Markov random field (Kindermann et al., 1980), each independent run being an observation and each region being a variable, to form a probabilistic undirected graphical model. This undirected graphical model formulates a probabilistic association between regions working together as a part of the same network to explain the experimental condition. We defined *functional connection probability* (FCP) between two regions, implementing the association factor for the undirected graphical model, by the joint probability of both regions *i* and *j* being important in separating the experimental conditions (*P*(*r_i_* ⋂ *r_j_*)). This probability can be obtained by the fraction of runs in which both regions *i* and *j* were *in-bag* and important (non-zero coefficient). A large FCP between two regions indicates that they are likely to be involved together in explaining the discriminative pattern of the response.

One should note that the standard association measures frequently used in Markov random fields, such as marginal correlation, sparse inverse covariance, and Kendall’s τ, are not sensitive to the means of association between regions in the interest of this study. As an example, these factors may indicate a high level of association between two regions that are highly correlated, but not necessarily important (low ISL) in most runs. This, however, is not desirable as we seek a factor that formulates the joint importance in explaining the experimental conditions.

FCP provides a measure of association (dependence) between regions being important for explaining the experimental conditions. Based on the Kolmogorov definition of the conditional probability, *P*(*r_i_* ⋂ *r_j_*) = *P*(*r_j_*)*P*(*r_i_|r_j_*), where *P*(*r_i_|r_j_*) is the conditional probability of a region *i* being important given that the region *j* is important. If region *i* and region *j* are independent then *P*(*r_i_* ⋂ *r_j_*) = *P*(*r_i_*)*P*(*r_j_*), which is smaller than each of *P*(*r_j_*) or *P*(*r_i_*), hence smaller than the ISL of *i* or *j*. When the importance of the two regions is dependent, however, the knowledge about the importance of one provides information about the importance of other, formulated in the term *P*(*r_i_|r_j_*).

The undirected graphical model is defined by a set of nodes, regions in this study, and vertices, the FCP between each two regions. The degree-related parameters of each node in this graph (e.g., node degree, degree centrality; Bullmore et al., 2012; Vecchino et al., 2017) indicate to what extent that node is required to function with other nodes to separate experimental conditions successfully. In the context of brain imaging and this study, this analysis reveals which brain regions are most likely to function together to describe the response patterns. The nodes with largest degree centrality, so called *hubs*, indicate the regions which are engaged the most with other regions to describe the response patterns. The decision points for each step of analysis are summarized in Figure 1.

**Figure 1.**
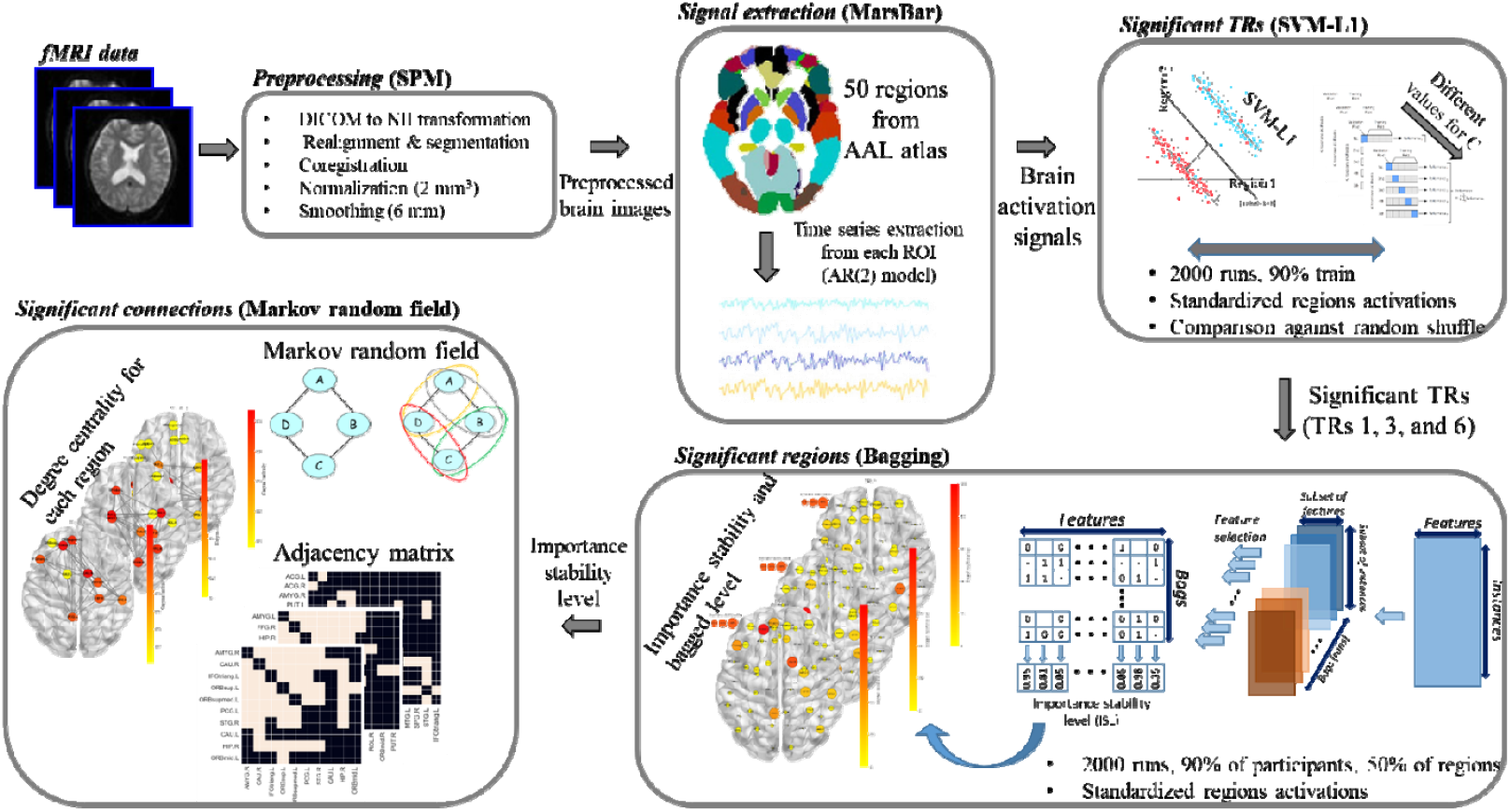
A flowchart summarizing decision made in each step of the analysis. Regions were selected from AAL atlas and weighted mean values were derived from each region of interest using Marsbar. SVM-L1 then applied on the brain signals. Importance stability level was used to identify regions that are important in distinguishing experimental conditions. We further used ISL to establish undirected graphical model to identify highly connected nodes (hubs) with high degree of centrality important in differentiating experimental conditions.

### Why SVM-L1?

While many studies have used the multivoxel pattern analysis (MVPA) to model classes that can distinguish between experimental conditions, this method has some limitations, in particular the curse of dimensionality and the issue of small sample size and a large feature size. A recent study (Jollans et al., 2019) has shown that, in the context of neuroimaging and MVPA, a sample size of over 400 is usually required to detect a small to moderate effect size when the number of features is large. To avoid this issue, we chose to summarize signals at the regional level, provided by MarsBar (see section “ Signal extraction” for details), over MVPA to reduce the impact of large feature size and small sample size in this study (Jollans et al., 2019). We used SVM for classification, a multivariate binary classification algorithm, that finds a hyperplane that best separates the classes. In comparison with other classification methods such as Logistic Regression and Partial Least Squares, SVM prioritises the hyperplane which has maximum distance from the instances in each class, minimizing the empirical risk. This approach presumably decreases the classification error on unseen instances (Cortes et al., 1995). L1 regularization in SVM leads to sparsity in the coefficients of the separating hyperplane, enabling the use of the coefficients as a measure for the importance of independent variables. This, of course, assumes that the independent identically distributed (*i.i.d*) assumption holds, which makes standardization (usually standardizing the variables to have a mean of zero and standard deviation of 1.0) essential for this conclusion to be accurate (Weston et al., 2001). In comparison with univariate methods such as ANOVA, SVM-L1 considers all independent variables at the same time, forming a sparse linear combination which best distinguishes the classes. This enables patterns to be described that involve multiple independent variables functioning together in a network. In the context of brain analysis, the classifier establishes a relationship between regions and conditions (Pereira et al., 2009). This means that the classifier assumes a dependency between brain responses (region activations) and conditions and formulates that relationship to be used for broader samples. SVM-L1 provides a set of sparse coefficients, forming a separating hyperplane (linear combination of the regions), while minimizes the empirical risk, enabling estimation of the importance of the regions while minimizing the error on the unseen samples.

### Behavioral analysis

We used two accuracy measures from signal detection theory: *False negative rate and False positive rate*. False negative rate is the number of incorrectly rejected responses divided by the number of statements that should have been rejected in each condition. This measures the tendency of participants towards rejecting statements without paying attention to the logical validity of the syllogism. False positive rate is the number of incorrectly accepted responses divided by the number of statements that should have been accepted in each condition. This measures the tendency of participants towards accepting statements without paying attention to the logical validity of the syllogism.

## Results

### Behavioural results

#### Endorsement rate

Endorsement rate, defined by the number of endorsed statements divided by the number of statements that should have been endorsed, differed significantly based on the believability of premises (*t*(31) = 3.24, *p* = 0.002, Cohen’s *d* = 0.83), with endorsement rate being higher for statements with believable premise (*M* = 1.11, *SD* = 0.23) than unbelievable premise (*M* = 0.95, *SD* = 0.16). However, no difference was found between believable and unbelievable conclusions, 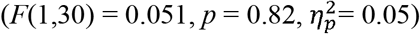.

#### Accuracy

It was found that the false negative rate is higher for statements with unbelievable premises (*M* = 0.14, *SD* = 0.14) than the statements with believable premises (*M* = 0.03, *SD* = 0.04; *t*(31) = 3.97, *p* = 0.0002, Cohen’s *d* = 1.07). In contrast, false positive rate was significantly higher for statements with believable premises (*M* = 0.25, *SD* = 0.24) than the statements with unbelievable premises (*M* = 0.12, *SD* = 0.14; *t*(31) = 2.5, *p* < 0.016, Cohen’s *d* = 0.66).

### Imaging results

#### Modelling the discriminative pattern of conditions

At each run for over 2000 runs, 90% of the participants were selected randomly for training purposes and the rest were left for testing. An SVM-L1 was trained on the training set with the original class labels and another SVM-L1 was trained on the training set when the class labels were shuffled randomly. The accuracy of the two models was calculated on the test set of that run. After 2000 runs, the accuracies of the models were compared using a *t-test* for each TR separately. As shown in Table 2, there is a significant difference between the original and the shuffled labels in TRs 1, 2, 3, 4, 5, 6, 10, 11, 17, indicating that the model can distinguish the original classes significantly better than chance in those TRs (all *p* < 0.001). As the most responses were recorded within the first 6 TRs, the rest of our analyses focused on TRs from 1 to 6 only. Further, a moderate to large effect size, measured by Cohens’ *d* larger than 0.5, was observed in TRs 1, 3, and 6.

**Table 2.**
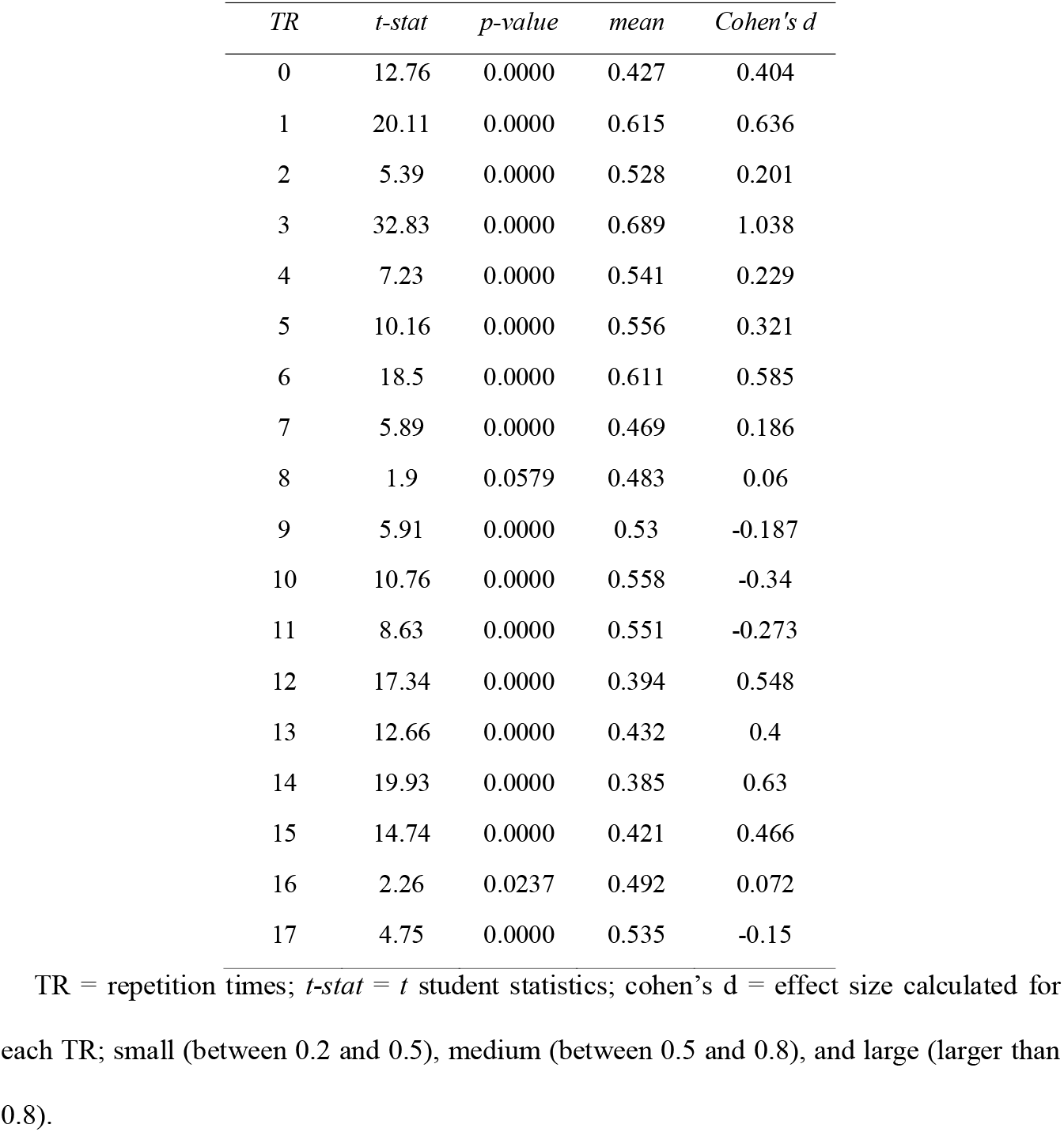
Inferential statistics of model performance for the 90% training dataset in 2000 runs compared with the shuffled data. The mean column provides the mean of accuracy measure in the test examples with no shuffling.

| *TR* | *t-stat* | *p-value* | *mean* | *Cohen's d* |
| --- | --- | --- | --- | --- |
| 0 | 12.76 | 0.0000 | 0.427 | 0.404 |
| 1 | 20.11 | 0.0000 | 0.615 | 0.636 |
| 2 | 5.39 | 0.0000 | 0.528 | 0.201 |
| 3 | 32.83 | 0.0000 | 0.689 | 1.038 |
| 4 | 7.23 | 0.0000 | 0.541 | 0.229 |
| 5 | 10.16 | 0.0000 | 0.556 | 0.321 |
| 6 | 18.5 | 0.0000 | 0.611 | 0.585 |
| 7 | 5.89 | 0.0000 | 0.469 | 0.186 |
| 8 | 1.9 | 0.0579 | 0.483 | 0.06 |
| 9 | 5.91 | 0.0000 | 0.53 | -0.187 |
| 10 | 10.76 | 0.0000 | 0.558 | -0.34 |
| 11 | 8.63 | 0.0000 | 0.551 | -0.273 |
| 12 | 17.34 | 0.0000 | 0.394 | 0.548 |
| 13 | 12.66 | 0.0000 | 0.432 | 0.4 |
| 14 | 19.93 | 0.0000 | 0.385 | 0.63 |
| 15 | 14.74 | 0.0000 | 0.421 | 0.466 |
| 16 | 2.26 | 0.0237 | 0.492 | 0.072 |
| 17 | 4.75 | 0.0000 | 0.535 | -0.15 |

TR = repetition times; *t-stat* = *t* student statistics; cohen's d = effect size calculated for each TR; small (between 0.2 and 0.5), medium (between 0.5 and 0.8), and large (larger than 0.8).

#### Importance stability factor (ISL) and bagged importance level (BIL) of regions

We calculated the ISL factor for TRs 1 to 6 as they were among TRs in which the machine learning method could distinguish between classes significantly better than chance. The ISL and BIL values for regions are shown in Figure 2. A full list of regions with their importance stability factors above 0.95 for each TR is presented in the GitHub link provided for this study. Among those TRs, TR 3 seems to have a large effect size (Cohen’s *d* larger than 0.8). Regions such as left amygdala, right hippocampus, left putamen, and left superior frontal gyrus were identified as significant (ISL>0.95) in this TR for differentiating between believable and unbelievable load.

One of the regions found to be important in differentiating the conditions was right insula in TRs 2, 3, and 4 (ISL 0.97, 0.99, and 0.99, respectively). Our results indicated the importance of subcortical areas such as left putamen in TRs 1, 2, and 3 (ISL 0.98, 0.96, and 1.00, respectively), right caudate in TR 6 (ISL 1.00), and right hippocampus in TRs 3 and 6 (ISL 1.00, 0.99, respectively) during the task. Another interesting region that was found to be involved in differentiating between our experimental conditions is the amygdala (right: TRs 1, 5, and 6, ISL 0.95, 0.97, and 0.96 respectively; left: TR 3, ISL 0.99) throughout the decision-making stage.

**Figure 2.**
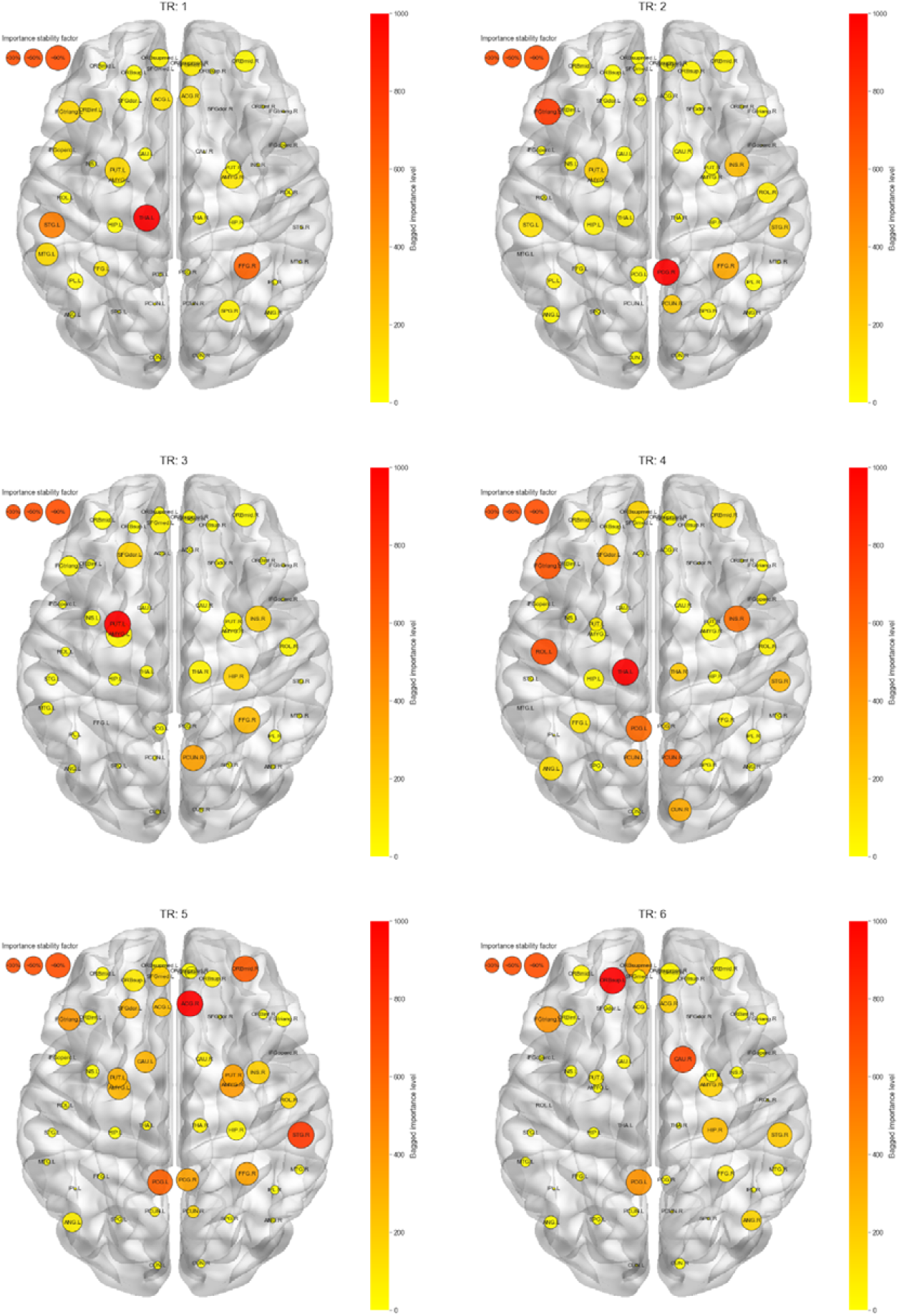
Importance of Regions. Regional importance in distinguishing experimental conditions derived form support vector machine (SVM-L1) are presented for each region, separated by repetition time (TR). Sizes indicate the importance stability factor (the larger the more likely to be important), colors indicate bagged importance level (the darker the more important, normalized from 0 to 1000).

#### Functional connection probability (FCP)

We next sought to determine what patterns of connectivity, defined by the joint probability of two regions being important, between regions of interest distinguish our experimental conditions in separate TRs (Figure 3). We focused on TRs 1, 3, and 6 for which a moderate to large effect size in distinguising experimental conditions was achieved. In those TRs, we only focused on region pairs with FCF > 0.95, i.e., the pair of regions which were important with a joint probablity larger than 95%. Finally, for each of these three TRs, we focused our analyses on the hubs, the nodes with highest degree centrality depicted in Figure 5.

**Figure 3.**
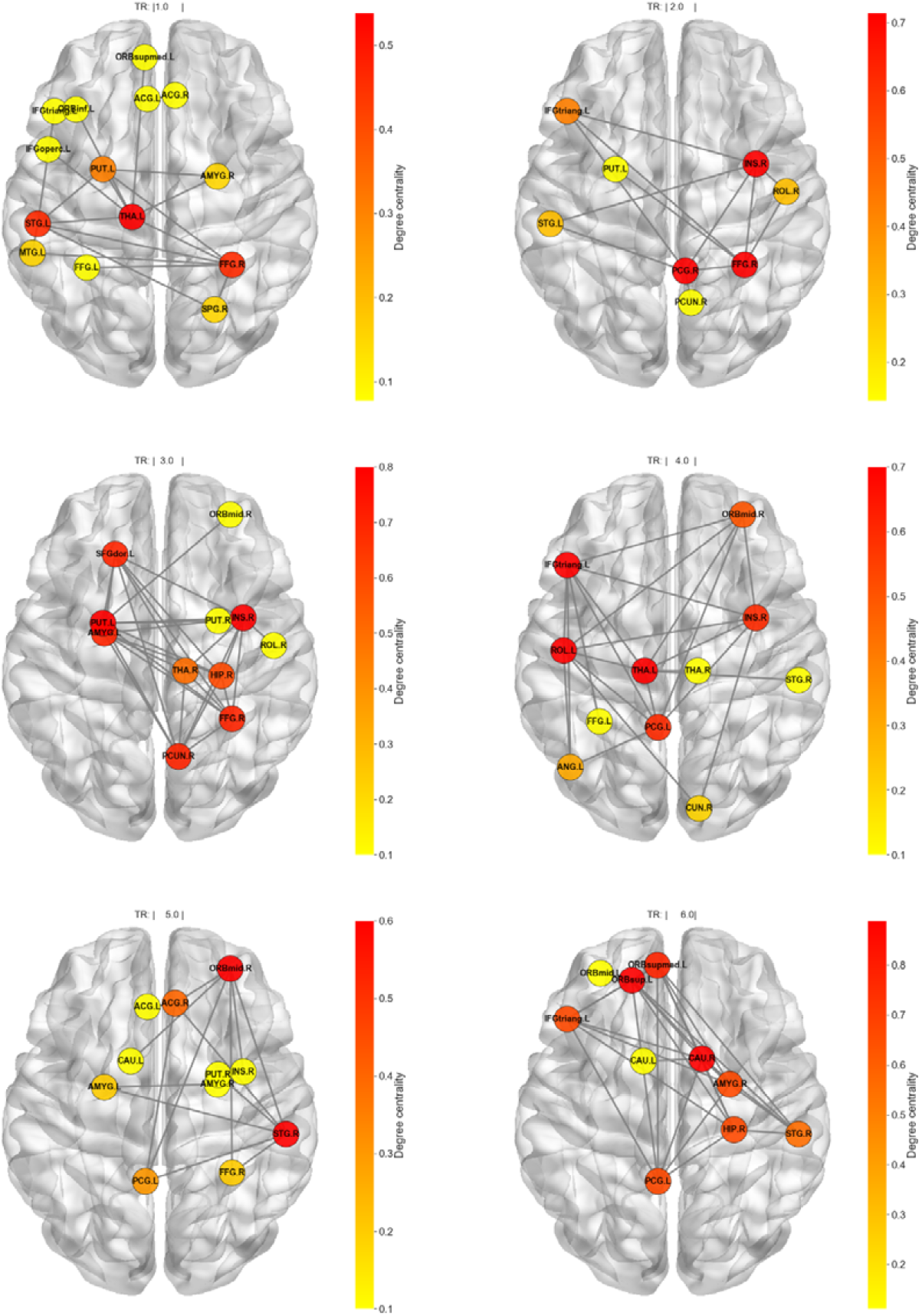
Undirected graphical model of functional connection of regions. The regions (nodes) with functional connection factor larger than 0.95 have been shown in the graph. Colors of the nodes represents the “degree centrality” (the lager, the more nodes that node is connected to).

Several novel findings have emerged from the connectivity analyses at these TRs. At TR 1 with moderate effect size (Cohen’s *d* of 0.63), thalamus was a highly connected node (degree centrality of 0.54) and was connected to regions such as fusiform gyrus, superior temporal gyrus, and putamen. At TR 3 with a large effect size (Cohen’s *d* of 1.03), insula and putamen had the largest degree centrality (degree centrality of 0.8). Insula was connected to the amygdala, hippocampus, putamen, fusiform gyrus, precuneus, and superior frontal gyrus. Amygdala and hippocampus were also directly connected. The putamen was connected to the amygdala, insula, hippocampus, superior frontla gyrus, precuneus, and fusiform gyrus. At TR 6 with effect size of 0.58, caudate and orbitofrontal cortex were the most connected regions (degree centrality of 0.89). The caudate was connected to regions such as the hippocampus, amygdala, orbitofrontal cortex and inferior frontal gyrus. Orbitofrontal cortex was also connected to caudate, amygdala, posterior cingulate gyrus, hippocampus, superior frontal gyrus and inferior frontal gyrus.

**Figure 4.**
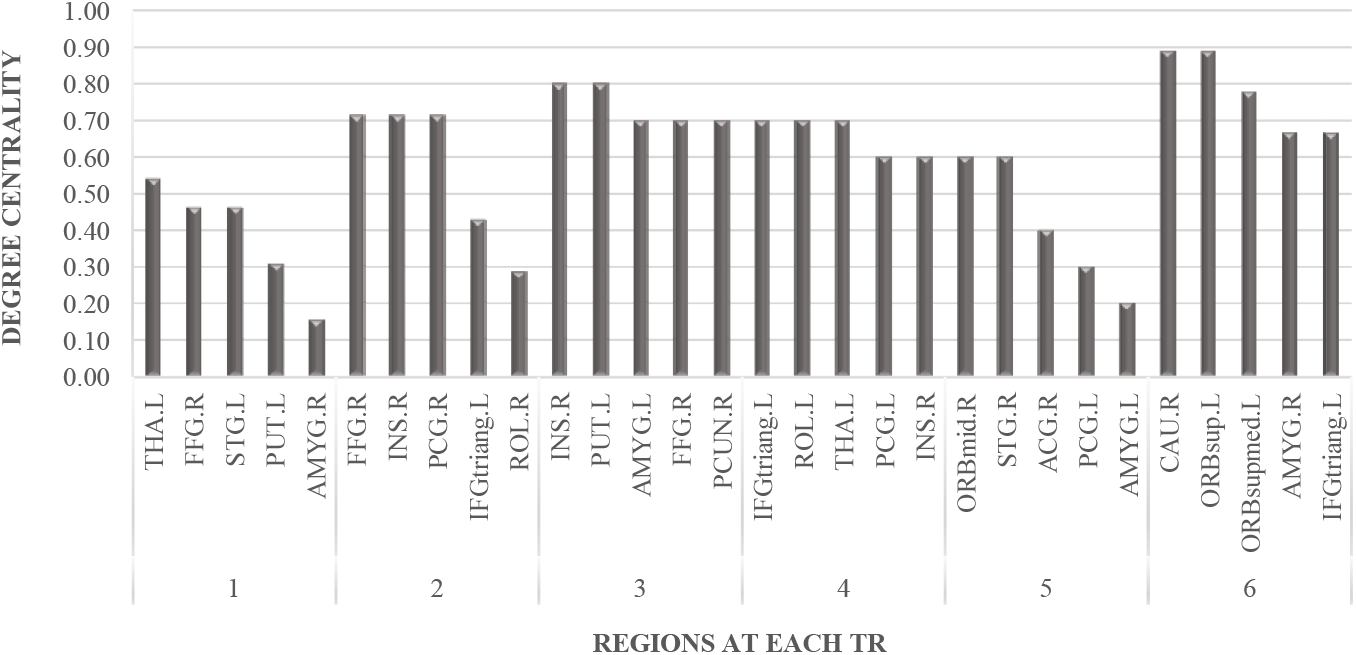
Degree centrality of regions at repetition time 1 to 6. The degree centrality of the top 5 regions have been reported across repetition time (TR)1 to 6. Functional connection factor has been thresholded at 0.95 for the calculation of the degree centrality. Abbreviations: TR = repetition time; AMY = amygdala; FFG = fusiform face gyrus; ORBinf = inferior orbitofrontal cortex; PUT = putamen; STG = superior temporal gyrus; THAL = thalamus; INS = insula; PCG = posterior cingulate gyrus; HIP = hippocampus; PCUN = precuneus; SFGdor = dorsal superior frontal gyrus; IFGtriang = Triangular inferior frontal gyrus; CAU = caudate; ORBsupmed = superior medial orbitofrontal gyrus.

## Discussion

Despite the number of studies that have investigated belief-logic bias in the literature, our understanding of the neural correlates underlying interaction between belief and logic remains limited. Importantly, there is a gap in the literature relating to the impact of the content of premises versus the content of conclusions on logical decision making. Given the importance of assumptions in our daily decision making, this study was set up to clarify these gaps in the literature using a novel analytical machine learning approach. Our behavioral results support the impact of belief load of premises, with participants favoring their own belief over the logical soundness of the statements. Using a regularized linear classifier, we were able to differentiate between believable and unbelievable assumptions given regional brain activity patterns. By using a bootstrap aggregation strategy, we found that the insula, hippocampus, amygdala, striatum, and inferior frontal gyrus are all important in distinguishing believable from unbelievable content before making a decision. We further introduced a measure for an association between regions, defined by the probability of pairs of regions functioning together. This association network established an undirected graphical model that revealed that connectivity anchored around the insula, amygdala, striatum, and the inferior frontal is critical for distinguishing between experimental conditions. Our task design and analytical approach allow us to speculate about cognitive mechanisms that underlie making a logical decision.

In line with our hypothesis, we anticipated that syllogisms with different belief loads would elicit emotional responses due to the manipulation of belief content. Although our task did not explicitly manipulate the emotional content of syllogisms, it is reasonable to assume that content-logic conflict elicits emotional responses, even implicitly, towards the syllogisms. Our results revealed that the content of assumptions, irrespective of emotional content, could interfere with the currently-held belief systems of decision-makers, which could elicit emotional responses in them. Previous studies have also suggested that decision making relies on a complex interaction between cognitive and emotional systems (Gupta et al., 2009). The importance of the amygdala in processing salience and ambiguous and emotionally-laden stimuli is also well established (Buhle et al., 2014; Urry et al., 2006; Ziaei et al., 2016), all of which are important processes involved in our logical reasoning task. Finally, another important function during reasoning is the ability to control emotional responses to logically decide and respond to the task. This might in turn activate the amygdala and prefrontal areas’ connections. The role of the amygdala is in accordance with the emotion regulation literature highlighting top-down regulatory control from prefrontal areas when personal emotions are required to be suppressed (Buhle et al., 2014; Ochsner et al., 2005). This pattern of connectivity seems to be important when making a logical decision. That is, our results resemble a regulatory pattern of connectivity between the inferior frontal gyrus, orbitofrontal cortex, and amygdala that are important in distinguishing believable from unbelievable inferences. Further studies are required to investigate the direction of interaction between limbic and frontal areas during the logical reasoning task.

During the logical reasoning task, integration of currently-held beliefs with externally given information in the syllogisms is warranted. No matter the outcome of this integration, the content of syllogisms activates the currently-held beliefs in reasoners. This function possibly engages the insula, a critical region important in integrating internal and external information to guide behaviour (Sridharan et al., 2008). The insula is implicated in a variety of cognitive and affective tasks, from cognitive control and attention orientation to emotional responses and empathy, making this region a candidate for detecting salient information (Menon, 2015; Menon et al., 2010). The engagement of insula during our task is not surprising given that integration of internal and external stimuli is needed to perform the task and thus the insula acts as a “switchboard” between the two worlds. Alternatively, due to the inconsistency between belief content and currently-held beliefs, a salient response might have been triggered engaging the insula. Salience processing is thought to play a role in decision making in emotionally driven situations (Eimontaite et al., 2019). Having said this, more research is needed to fully investigate the link between salience rating and participants ‘ emotional responses towards syllogisms that include different belief content.

Confirming our expectations and in line with previous studies (Goel et al., 2004b; Ziaei et al., 2020a), the hippocampus was engaged at a relatively early stage of the task to possibly retrieve currently-held beliefs stored in memory to compare with given assumptions. The amygdala-hippocampal connectivity concords with the memory-modulation hypothesis that the arousal level induced by items is a critical factor for memory (Mather et al., 2011). Extending previous work (Preston et al., 2013; Ziaei et al., 2020a), we demonstrated that memory processing and retrieval of semantic information played a critical role during the logical reasoning task engaging the hippocampus in connection with the prefrontal areas and the insula. We speculate that this pattern of activity might suggest that retrieving currently-held beliefs from memory is required during the reasoning task. There is still a need for further investigation to highlight the exact link between memory and syllogistic reasoning ability in more detail.

One other important component of a decision making is planning of a response (Melrose et al., 2007), before making a response, which is expected to elicit striatum activation. In line with our results, a previous study of intense training in logical reasoning reported a strengthening of prefrontal-parietal and parietal-striatal connections following training, possibly through dopaminergic inputs (Ashby et al., 2010), supporting the importance of the striatum in reasoning (Mackey et al., 2013). A link between dopaminergic neurotransmission in the striatum and salience detection in decision-making tasks has also been reported previously (Esslinger et al., 2013; Rausch et al., 2014). Only a few studies, however, have reported activity of the putamen specifically during the premise integration stage and during reasoning (Eimontaite et al., 2019; Reverberi et al., 2012). Additionally, some studies have confirmed functional and structural connectivity between the putamen and prefrontal areas (Di Martino et al., 2008) and the insula (Postuma et al., 2005). Altogether, cognitive mechanisms involved in our task suggest that activities and connections between prefrontal and subcortical areas are essential in transitioning from planning and integrating assumptions to forming a logical decision and implementing a logical response.

In line with our expectation, the inferior frontal gyrus was activated during the task possibly to inhibit unnecessary responses and assumptions. This region has received lots of attention in the literature highlighting the role of inhibitory control during logical reasoning (Ziaei et al., 2020b). There is mounting evidence that the inferior frontal gyrus plays a key role during both emotional and non-emotional reasoning tasks (Prado et al., 2011; Rotello et al., 2014; Tsujii et al., 2011). In the current study, the role of IFG has been further delineated by its connection with the striatum, limbic, and salience hubs at a later stage of reasoning confirming our predictions about the importance of inhibitory control over currently-held beliefs before making a decision.

Our imaging and analytical method allow us to expand previous neurocognitive models of reasoning and propose a link between neural and cognitive processes involved in different stages of reasoning. It has to be noted that although we did not measure the cognitive processes in each stage of the reasoning separately, our high temporal resolution data allows us to investigate the different neural and cognitive mechanisms involved in each stage of reasoning prior to making a logical decision. Here, we integrate previous dominant theories, such as the Mental Models (Johnson-Laird, 2010) and the neurocognitive model of reasoning (Knauff, 2009), with our results to offer a modification for the multi-stage process, that relies on several cognitive and neural processes.

While some empirical studies have initiated the investigation of brain regions underlying cognitive processes of reasoning (Fangmeier et al., 2006; Reverberi et al., 2012a), the brain regions related to the underlying cognitive functions have not been fully mapped out. In the earlier stage of reasoning (TR 1-2), it is speculated that a mental image of assumptions will be constructed, activating regions such as the thalamus and visual and parietal areas. During the middle stage (TR 3-4), content-logic conflict appears to trigger the retrieval of currently-held beliefs from memory and elicit emotional responses, activating regions such as the hippocampus and amygdala, mediating memory and affective response respectively. At this stage, integrating internal and external information is required activating the insula as a functional ‘switchboard’. Subsequently, inhibition of retrieved beliefs and emotional responses are necessary for logical decision making, and prefrontal regions are engaged to validate the final decision during the late stage of the decision-making process (TR 5-6; Figure 5).

Unlike most previous studies, our approach facilitates the detection of multiple regions that are simultaneously involved in reasoning at different time points. Future behavioral and neuroimaging studies are needed to confirm and extend our proposed model using reasoning tasks that vary in their emotional and conflict contents. Additionally, given that our analyses aimed to represent important regions at different time points, future studies are needed to examine the chronological dependencies of the processes at different stages of reasoning.

**Figure 5.**
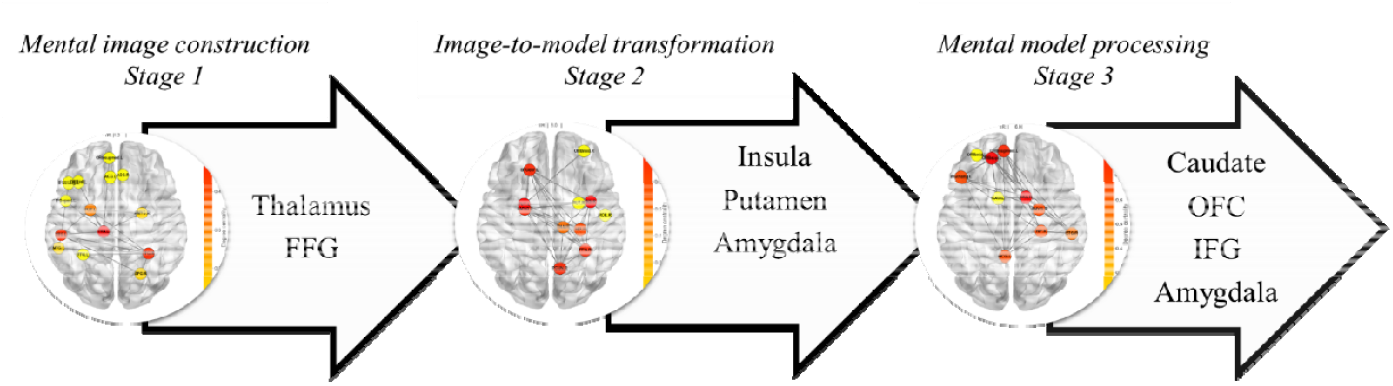
A schematic brain-based model of decision-making process involved in predicting belief load. Regions such as thalamus and fusiform face areas were involved during the initial stage of reasoning (mental image construction stage). Regions such as insula, amygdala, striatum and hippocampus were primarily engaged during the middle stage of reasoning (around repetition time (TR) 3 or comprehension stage) to possibly link between internal and external information for making a decision. Regions such as Inferior frontal gyrus and orbitofrontal were among the regions that were mostly activated and connected to other areas during the final stage (TR 6; validation stage) in which the final decision was being validated. Abbreviation: FFG = fusiform face gyrus; OFC = orbitofrontal cortex; IFG = inferior frontal gyrus.

## Conclusion

We have shown that the belief load of premises affects logical decision making during the conclusion stage of the decision-making process. Multiple brain areas were found to be important in distinguishing belief loads manipulated before a logical decision being made. By employing a machine learning method, we were able to further identify *connections* between different brain areas such as IFG, insula, and limbic areas that play a critical role in discriminating between believable and unbelievable premises. Our results shed light on the interaction between multiple brain areas, and the underlying cognitive mechanisms involved in belief bias. Our results offer a closer mapping between brain and cognitive functions involved in reasoning based on the believability load of assumptions, allowing us to extend existing brain-based models of logical decision making.

## Acknowledgment

The authors would like to thank our participants for their time and acknowledge the practical support provided by the imaging staff at the Centre for Advanced Imaging. We would also like to thank Ms. Ashley York for her help with the data collection. This work was supported by funding from the Australian Research Council Science of Learning Special Research Initiative (SR120300015) to DCR.

Authors declare no conflict of interest.

## Notes

### Competing Interest Statement

The authors have declared no competing interest.

